# Reduction of *Bacillus cereus* Biofilms on Stainless Steel Surfaces by *Pestalotiopsis* sp. Culture Crude Extract

**DOI:** 10.1101/628990

**Authors:** Rener S. De Jesus, Gina R. Dedeles

## Abstract

*Bacillus cereus* is capable of forming biofilms on many types of substrata, including stainless steel surfaces. Thus, the presence of *B. cereus* biofilms in food production facilities is inevitable, and possible food contamination could occur, resulting in product spoilage and deterioration, which would eventually lead to product rejection and financial loss. *Pestalotiopsis* spp. have been shown to produce a wide range of novel secondary metabolites that have potential medicinal, industrial, and agricultural applications. In this study, *B. cereus* biofilms were grown on milk pre-soiled stainless steel coupons (20 × 5 mm) for 48 hr at 30°C in nutrient-depleted medium. After incubation, the biofilms on the coupons were immersed for 15 min in different concentrations (25, 50, or 100 mg/mL) of *Pestalotiopsis* sp. culture extract at 25°C or 50°C. The results showed that the 48-hr old *B. cereus* biofilms were significantly reduced (p < 0.05) following immersion in all concentrations of extract. Specifically, the 100-mg/mL concentration resulted in 0.18- to 2-log_10_ and 0.48- to 2.28-log_10_ reductions at 25°C and 50°C, respectively. Secondary metabolites were detected in *Pestalotiopsis* sp. culture crude extract using thin layer chromatography, indicating that these metabolites have antibiofilm activity. Additionally, fluorescence micrographs displayed decreasing light intensities emitted by the fluorescent dye ethidium bromide at 605 nm, as was also evident from the interactive 3-D surface plots generated using ImageJ 1.52a software, which indicated that the extracellular polymeric substances in the biofilms were disrupted. The current study concluded that secondary metabolites from the extract of *Pestalotiopsis* sp. have antibiofilm activity and contain possible bioactive agents with activity against many varieties of microbial biofilms.

## 1. Introduction

*Bacillus cereus* is frequently present in food production environments because of its adhesive and resistant endospores (Stenfors Arnesen et al., 2008). Additionally, its vegetative cells are capable of producing various toxins (Beecher & MacMillan, 1991; Lund & Granum, 1996) that, upon consumption of contaminated foods, can cause diarrheal and emetic types of food poisoning. This bacterium is a gram-positive, aerobic or facultatively anaerobic, spore-forming rod that is phenotypically and genetically related to other *Bacillus* species, especially *Bacillus anthracis* (Ash et al., 1991). Recently, an emerging *B. cereus* strain that causes anthrax-like disease was reported, and this strain carries *B. anthracis*-like plasmids that encode toxins and polyglutamate capsules (Brézillon et al., 2015). Aside from toxin production, *B. cereus* can also cause product spoilage due to its ability to produce hydrolytic extracellular enzymes, such as proteases (Kim et al., 2001) and lipases (Chen et al., 2007), which can greatly affect dairy products. A number of researchers have isolated *B. cereus* from milk filling machines (Eneroth et al., 2001), dairy processing lines (Faille et al., 2001), and silo tanks (Svensson et al., 2004); therefore, these areas are possible sources of milk product contamination. Furthermore, *B. cereus* has also been isolated and detected in ready-to-eat foods (Zhang et al., 2016), raw meat products (Tewari et al., 2015) and fried rice dishes (Chang et al., 2011). Since the spores of *B. cereus* are persistent in food production areas, vegetative cells could grow from the persistent spores and eventually lead to biofilm formation.

Biofilms are microbial communities that are irreversibly attached to surfaces or to each other, are embedded in a matrix of extracellular polymeric substances (EPS) that the bacteria produced, and exhibit altered phenotypes with respect to growth rate and gene transcription (Donlan & Costerton, 2002). Microbial biofilms containing *B. cereus* are of great concern in the food industry, such as in fresh products and in poultry, dairy and raw meat processing facilities, and are an impending source of recurrent cross-contamination and postprocessing contamination of finished products, which results in spoilage and foodborne illnesses (Rajkovic et al., 2008). Generally, microbial biofilms are resistant to antibiotics and disinfectants because of their self-produced EPS that acts as “protective casing.” Furthermore, EPS helps maintain intercellular communication (e.g., horizontal gene transfer) between biofilm cells, making survival possible. Because of these characteristics, biofilm eradication and control programs are one of the challenges facing the food industry today. Various chemical disinfectants, such as hypochlorites, phenolic substances and quaternary ammonium compounds (also known as Quats) are presently used by many food production industries to eliminate spoilage and pathogenic microorganisms and to prevent the formation of microbial biofilms. However, because of their complex biological mechanisms and EPS, most microbial biofilms are able to tolerate these chemical disinfectants and, hence, proliferate in food production systems (Bayoumi et al., 2012; Yang et al., 2016; Møretrø et al., 2017).

The genus *Pestalotiopsis* is a group of ascomycete fungi widely distributed in tropical and temperate regions of the world, and members of this genus can be isolated as plant endophytes. Numerous studies have been published that document the many novel secondary metabolites with medical, agricultural, and industrial uses that are produced by various species of this genus (Hwang et al., 2015; Li et al., 2015; Sharma et al., 2016), demonstrating *Pestalotiopsis* to be a rich source for bioprospecting. To date, various endophytic fungal species have been studied for secondary metabolites that exhibit antibiofilm activity. Examples of these include the new phenyl derivatives isolated from an endophytic fungus, *Aspergillus flavipes*, which were shown to penetrate through the biofilm matrix and kill live cells inside mature *Staphylococcus aureus* biofilms (Bai et al., 2014). Other examples include the bioactive specialized metabolites extracted from a mangrove-derived endophytic fungus, *Eurotium chevalieri*, which were shown to inhibit biofilms of *S. aureus* and *Escherichia coli* (Zin et al., 2017). Presently, there have been no specific studies conducted on the antibiofilm activities of secondary metabolites extracted from *Pestalotiopsis* sp. Therefore, this study aimed to reduce *B. cereus* biofilms grown on stainless steel surfaces using a *Pestalotiopsis* sp. culture crude extract, which could be a promising antibiofilm agent and possible alternative method for biofilm elimination in the food industry.

## 2. Materials and methods

### 2.1 *B. cereus* and *Pestalotiopsis* sp. test isolates

All *B. cereus* test organisms used in this study were isolated from four different soil samples and five samples of infant formula milk powder by the spread plate method using *B. cereus* agar (BCA). The isolates were identified based on their bacterial morphology, colony morphology, and conventional biochemical characteristics. The *Pestalotiopsis* sp. was isolated from a *Mangifera indica* leaf sample by the surface sterilization method (Schulz et al., 1993) using quarter-strength potato dextrose agar (PDA). The fungal colony was identified according to its conidial morphology and culture characteristics. The axenic *B. cereus* isolates were stored in cryovials containing trypticase soy broth (TSB) with 15% glycerol. The *Pestalotiopsis* isolate was grown on a PDA slant and was then overlaid with sterile mineral oil. The isolates were deposited at the University of Santo Tomas Collection of Microbial Strains (UST-CMS).

### 2.2 Microtiter plate biofilm formation assay

All *B. cereus* isolates were subjected to microtiter plate (MTP) biofilm formation assays to select isolates for biofilm formation on stainless steel coupons. Following the method of Martinez-Medina et al. (2009), overnight cultures of *B. cereus* were grown in MTP wells containing 200 μL diluted (1:20 [v/v]) sterile TSB for 24 hr at 30°C. After incubation, the broth was removed and the wells were replenished with full-strength sterile TSB and incubated again at 30°C for 24 hr. The optical densities (ODs) of the wells were read at 630 nm using an absorbance microplate reader (SH-1000Lab, Corona Electric Co. Ltd., Japan), the broth from each well was removed, and the wells were then washed with 200 μL sterile phosphate buffered solution (PBS), pH 7.2. The MTP was air dried for 20 min and was then stained with 130 μL 1% crystal violet (CV) solution for 5 min. After staining, the CV solution was removed, the MTP wells were washed thrice with sterile distilled water, the remaining CV was then solubilized in 130 μL absolute ethanol, and the ODs were measured at 570 nm. The biofilm measurements were calculated using the following formula: *SBF*=(*AB*-*CW*)/*G*, in which *SBF* stands for specific biofilm formation, *AB* is the OD at 570 nm, *CW* is the OD at 570 nm of stained control wells containing only uninoculated medium (to eliminate nonspecific or abiotic absorbance), and *G* is the OD at 630 nm measuring the cell growth in broth (Niu & Gilbert, 2004). In each assay, staining and analysis was performed in triplicate for each isolate and for the control wells. The degree of biofilm production was classified according to three categories adapted from Martinez-Medina et al., 2009: weak (SBF<0.5), moderate (0.5 – 1.0), and strong (SBF>1.0).

### 2.3 *Pestalotiopsis* sp. culture crude extraction and thin layer chromatography

Using the method of Zhao et al. (2010) with modification, five agar plugs (1 × 1 cm^2^) from a 5-day old culture of *Pestalotiopsis* sp. grown on PDA were transferred into Erlenmeyer flasks containing 500 mL sterile potato dextrose broth (PDB). The flasks were incubated at 28°C with constant shaking at 150 rpm/min (Model 420 Series, Forma Scientific Orbital Shaker, Thermo Scientific, U.S.) for three weeks. After incubation, a total of 15 L fermentation broth was harvested and transferred into sterile Erlenmeyer flasks while passing through three layers of sterile cheesecloth to separate the mycelia and filtrates. The filtrates were filtered again using fluted filter paper, transferred into new sterile flasks and extracted three times with ethyl acetate at a sample:solvent ratio of 1:1 (v/v) using a separatory funnel. The solvent phase (top layer) was collected after each extraction, and the final extract was evaporated to dryness using a rotary evaporator (Eyela N-1200A, Tokyo Rikakikai Co. Ltd.) under reduced pressure at 40°C. The dried residues were collected and stored at 4°C for further analysis.

For thin layer chromatography (TLC), TLC plates (3 × 10 cm) (TLC Silica Gel 60 F_254_, Merck) were spotted by simply touching the end of a capillary tube containing fungal crude extract to the coated side of the TLC plates. The plates were then placed in 400 mL beakers containing solvent systems, which were then covered with a glass lid. The four solvent systems were (1) 9:1 chloroform/methanol, (2) 6:3 chloroform/acetonitrile, (3) 9:1 chloroform/ethyl acetate, and (4) 3:1 ethyl acetate/hexane. After the development of the spots, the TLC plates were viewed under UV lights (MinUVIS; Desaga, Heidelberg, Germany) at 254 and 366 nm. The spots were cut and placed on 24-hr old lawns of *B. cereus* isolates on trypticase soy agar (TSA) agar plates to determine the antibacterial activity.

### 2.3 *B. cereus* biofilms on stainless steel surfaces

#### 2.3.1 Preparation of spore suspensions from B. cereus isolates

*B. cereus* sporulation was induced by the addition of 10 g anhydrous manganese sulfate (MnSO_4_) per liter of nutrient broth (NB). Following the method of Elhariry (2011) with some modifications, the *B. cereus* isolates were inoculated into two Erlenmeyer flasks containing 100 mL NB-MnSO_4_ and incubated at 35°C for 5 days. The spores were harvested by transferring 5 mL broth culture into sterile centrifuge tubes and subsequently washing the pellet following repeated centrifugation (10 min at 800 x*g* and 22±2°C). After the supernatant was discarded, sterile distilled water was added to the pellet, and the pellet was heated using a water bath to 80°C for 10 min to kill the remaining vegetative cells. The tubes were then held for 5 min at room temperature and centrifuged again (15 min at 10,000 x*g* and 22±2°C). The spore pellets were resuspended in 10 mL sterile PBS, pH 7.2 and then stored at 4°C until use. The spore density in 10 mL sterile PBS was approximately 6 log_10_ spores per mL, as verified by the spread plating technique.

#### 2.3.2 Preparation of milk pre-soiled stainless steel coupons

Stainless steel coupons (SSCs) (20 × 5 mm; type 304L) were prepared by soaking for 1 hr in a 1:1 (v/v) mixture of ethanol and acetone to remove grease and then rinsing with deionized water. After rinsing, the coupons were autoclaved for 15 min at 121°C and dried overnight at 50°C in a laboratory oven (Memmert GmbH + Co. KG). The dried SSCs were submerged in low-fat, UHT-sterilized milk for 1 hr at ambient temperature and were subsequently rinsed with sterile deionized water before immediate use (Wijman et al., 2007). The milk was tested for commercial sterility, and no microbial growth was observed.

#### 2.3.3 Attachment of spores and formation of biofilms on SSCs

The coupons were immersed in *B. cereus* spore suspensions for 5 sec and then transferred into sterile vials positioned at an incline and containing 1 mL diluted TSB (1:20 [v/v]). Afterwards, the vials were incubated statically at 30°C for 24 hr. After incubation, the coupons were gently removed from the vials and rinsed with sterile water to remove unattached planktonic cells. The coupons were then transferred into new sterile vials containing 1 mL full-strength TSB and incubated again under the same conditions as described earlier. Blank controls were vials containing SSCs and TSB broth only. After incubation, the coupons were used for either immersion in the fungal extract or for CV staining to visualize the biofilms.

### 2.4. Reduction of biofilms on SSCs by *Pestalotiopsis* sp. culture crude extract

The *B. cereus* biofilms grown on milk pre-soiled SSCs were immersed for 15 min in 100, 50, or 25 mg/mL *Pestalotiopsis* sp. culture crude extract at two different temperatures: 25°C or 50°C. Each condition was tested in triplicate. A total of 27 SSCs for each *B. cereus* test isolate were tested. The blank control coupons were immersed in sterile water only (non-treated samples). After immersion, all SSCs were carefully rinsed with sterile water, transferred into centrifuge tubes with sterile PBS, pH 7.2, vigorously shaken, serially diluted with 0.1% peptone in water and plated on sterile Petri plates with molten TSA to enumerate the remaining viable *B. cereus* biofilm cells. After 48 hr of incubation at 30°C, the *B. cereus* colony-forming units (CFUs) that were visible on the plates were counted, and the mean was calculated and expressed in logarithmic form (Log_10_ CFU/coupon). The means and standard deviations were analyzed for statistical significance using t-tests and one-way analysis of variance (ANOVA).

### 2.5 Fluorescence microscopy and biofilm imaging

The *B. cereus* biofilms on SSCs before or after immersion in *Pestalotiopsis* sp. culture extract were stained with 1% ethidium bromide (EtBr) for 10 min. The coupons were rinsed carefully with sterile water, air dried and then viewed under a fluorescence microscope (Olympus BX43) with an attached camera (Olympus DP25) and UV illumination at 605 nm. The fluorescence micrographs were analyzed using the ImageJ (Image Processing and Analysis in Java) 1.52a software (http://imagej.nih.gov.ij). The SSC biofilms were viewed using a high power objective (HPO), and the micrographs used for ImageJ analysis were approximately 131 × 131 pixels in size. The software generated interactive 3-D surface plots, where the z-axis indicated the recorded intensities of each coordinate within the 131 × 131 pixels (the x- and y-axes).

## 3. Results

### 3.1 Production of *B. cereus* biofilms in MTPs

A total of 12 *B. cereus* test isolates were evaluated for their ability to produce biofilms by MTP biofilm formation assay (Table 1). Biofilms were visually evident to be forming on the walls of the MTP wells after only 24 hr of incubation at 30°C under static and nutrient-depleted conditions. After an additional 24 hr of incubation and replenishing of the MTP wells with full-strength sterile TSB, the biofilms became more visible and noticeably thicker. Based on the mean OD readings measured using the microplate reader, four isolates were identified as strong (S) biofilm producers and eight were moderate (M) biofilm producers (Table 1). The highest SBF value was 2.341±0.642, attained by isolate BC-MS010. None of the isolates were identified as weak (W) biofilm producers. The four isolates identified as S biofilm producers were selected for biofilm formation on SSCs and were isolates BC-SS001, BC-SS006, BC-MS003, and BC-MS010.

**Table 1.** Results of microtiter plate biofilm formation assay showing the optical density readings of *B. cereus* biofilms in microtiter plate wells and their assigned biofilm formation categories.

| Isolate Codes | Specific Biofilm Formation (SBF)<br>Mean Optical Density (OD) Readings | | | mean SBF ± SD | Biofilm Formation<br>Category (BFC) |
| --- | --- | --- | --- | --- | --- |
| Control | 1.060 | 1.049 | 1.028 | - | - |
| BC-SS001 | 1.893 | 1.110 | 1.557 | 1.520 ± 0.393 | S |
| BC-SS002 | 0.927 | 0.112 | 1.123 | 0.721 ± 0.536 | M |
| BC-SS004 | 1.107 | 0.200 | 0.890 | 0.731 ± 0.474 | M |
| BC-SS006 | 2.178 | 2.364 | 1.877 | 2.140 ± 0.246 | S |
| BC-SS007 | 0.701 | 0.368 | 1.008 | 0.692 ± 0.320 | M |
| BC-MS001 | 0.609 | 0.643 | 0.738 | 0.663 ± 0.067 | M |
| BC-MS003 | 0.938 | 1.238 | 0.894 | 1.023 ± 0.188 | S |
| BC-MS004 | 1.100 | 0.739 | 0.990 | 0.943 ± 0.185 | M |
| BC-MS005 | 1.015 | 0.813 | 0.875 | 0.901 ± 0.103 | M |
| BC-MS008 | 0.582 | 0.430 | 0.785 | 0.600 ± 0.180 | M |
| BC-MS009 | 0.643 | 0.942 | 0.930 | 0.838 ± 0.170 | M |
| BC-MS010 | 2.108 | 1.848 | 3.067 | 2.341 ± 0.642 | S |

**Note:** BFCs: **W**, weak biofilm producer; **M**, moderate biofilm producer; and **S**, strong biofilm producer. **SD**, standard deviation

### 3.2 Formation of *B. cereus* biofilms on SSCs

Spore suspensions from 5-d old cultures of BC-SS001, BC-SS006, BC-MS003, and BC-MS010 isolates were prepared and used for biofilm formation assays on milk pre-soiled sterile SSCs. The *B. cereus* biofilms on SSCs were clearly observable after a 48 hr incubation period at 30°C under static, nutrient-depleted conditions and formed preferentially at the air-liquid interface (ALI). However, biofilms also extended below the ALI, where O_2_ was at a low concentration or was completely absent; these biofilms were thinner and weaker than the ALI biofilms and were easily detached by simple rinsing. The *B. cereus* biofilms on the SSCs were then stained with CV solution to enable visualization, as shown in Figure 1.

**Figure 1.**
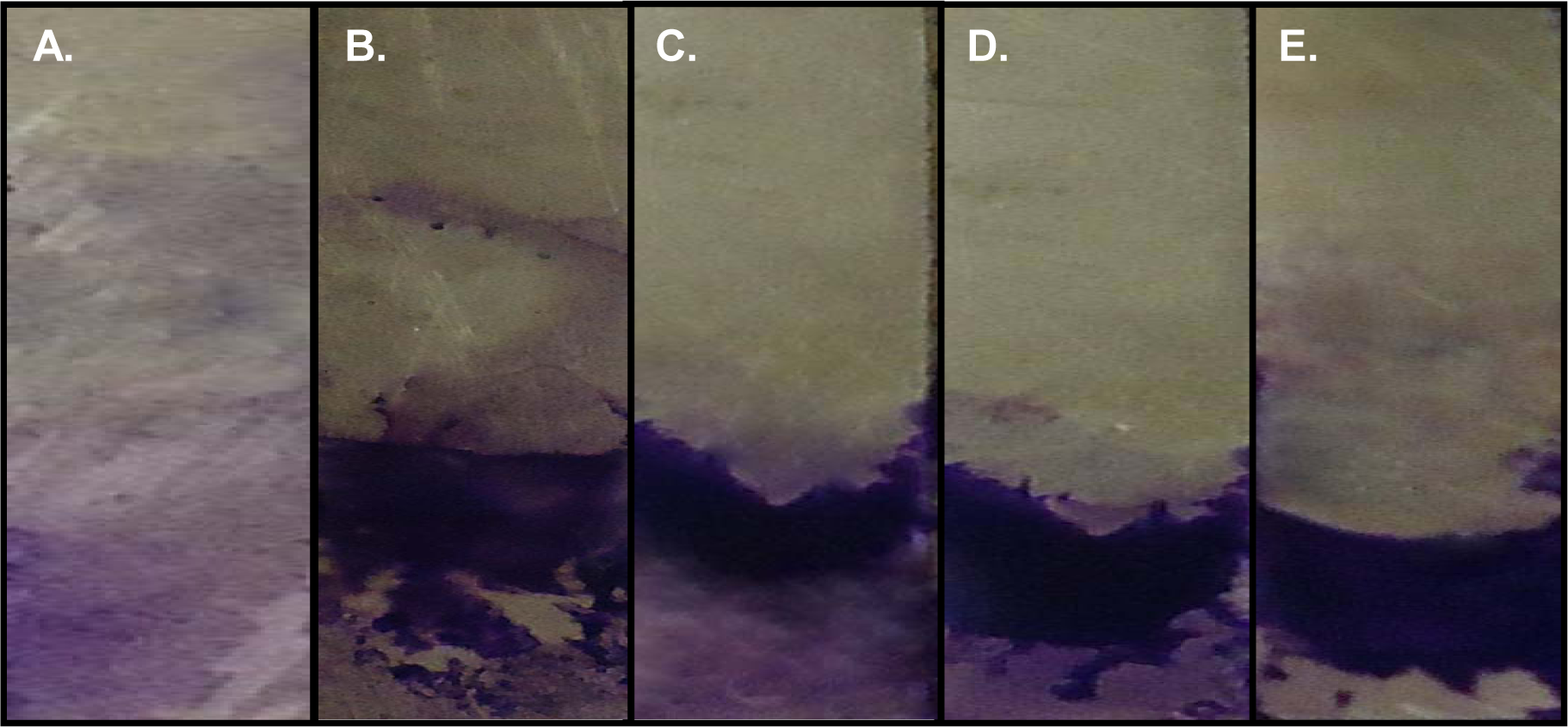
Staining with 1% crystal violet shows 48-hr old *B. cereus* biofilms formed at the ALI on milk pre-soiled SSCs. (**A**) Control; (**B**) BC-SS01; (**C**) BC-SS006; (**D**) BC-MS003; (**E**) BC-MS010.

### 3.3. Reduction of *B. cereus* biofilms by *Pestalotiopsis* sp. culture crude extract

The *B. cereus* biofilms on milk pre-soiled SSCs were immersed in 100, 50 or 25 mg/mL *Pestalotiopsis* sp. culture crude extract for 15 min at two different temperatures: 25°C or 50°C. After immersion in 25 mg/mL crude extract, the number of biofilm cells was significantly reduced (p < 0.05), resulting in 5.44 log_10_ CFU/coupon. When immersed in 50 mg/mL crude extract, 4.34 log_10_ CFU/coupon remained, while exposure to 100 mg/mL crude extract resulted in 3.62 log_10_ CFU/coupon (Figure 2). Hence, the crude extract at 25°C resulted in a 0.18- to 2-log_10_ reduction in *B. cereus* biofilm cells on SSCs. At 50°C, the number of biofilm cells was significantly reduced as well (p < 0.05), resulting in 5.15, 3.93, and 3.35 log_10_ CFU/coupon for the 25, 50, and 100 mg/mL concentrations, respectively. Based on these results, the *Pestalotiopsis* sp. culture crude extract at 50°C resulted in a 0.48- to 2.28-log_10_ reduction of *B. cereus* biofilm cells on the SSCs. The control (non-treated) biofilms had averages of 5.6 and 5.5 log_10_ CFU/coupon at 25°C and 50°C, respectively.

**Figure 2.**
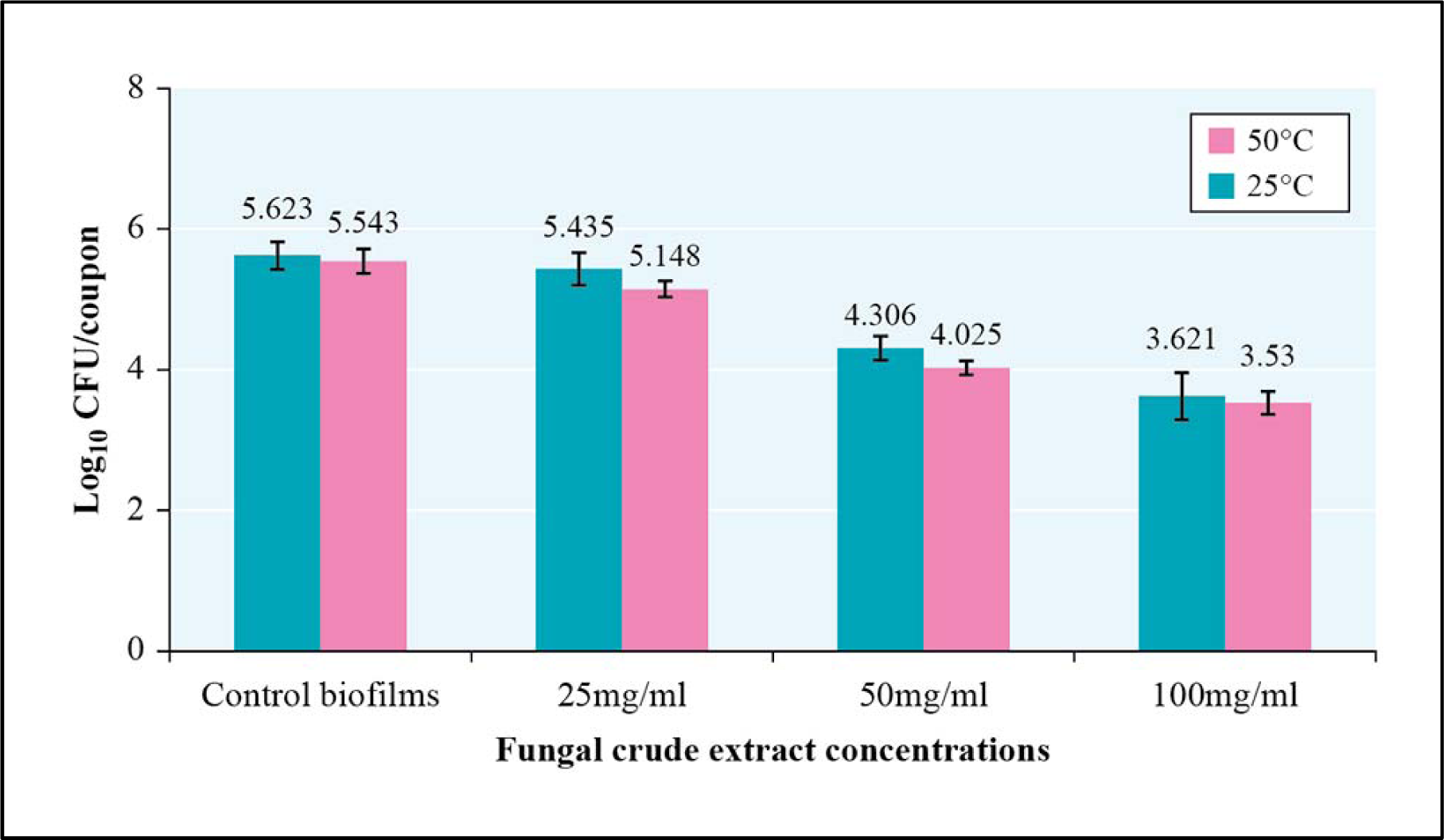
Reduction of *B. cereus* biofilms on milk pre-soiled SSCs after 15 min exposure to different concentrations of *Pestalotiopsis* sp. culture crude extract at 25°C or 50°C. The crude extract was tested at concentrations of 25, 50, and 100 mg/mL. Error bars indicate the standard deviation from the mean (n=4).

The crude extract was found to contain secondary metabolites as detected in TLC plates under UV lights, indicating that these metabolites might have activity against *B. cereus* biofilms on SSCs. Among the four solvent systems, the greatest number of secondary metabolites (9) was detected using 9:1 chloroform/ethyl acetate. However, these metabolites did not exhibit antibacterial activity against *B. cereus* on TSA agar plates. Therefore, the secondary metabolites in *Pestalotiopsis* sp. culture crude extract did not lyse the *B. cereus* vegetative cells but could disrupt the biofilm EPS on stainless steel surfaces.

### 3.4 Fluorescence micrographs of *B. cereus* biofilms on SSCs

Fluorescence micrographs of EtBr-stained *B. cereus* biofilms were analyzed for their EtBr-emitted light intensities (red-green-blue, or RGB values) using ImageJ 1.52a software, and interactive 3-D surface plots were then generated using the same program. As illustrated in Figure 3, the mean recorded emitted light intensities decreased drastically, suggesting that there was disruption of biofilm components after 15 min immersion in different concentrations of *Pestalotiopsis* sp. culture crude extract at 25°C and 50°C of BC-MS010 biofilms. This trend was further revealed by the 3-D surface plots of BC-MS010 biofilms, as shown in Figure 4. The same trends were also observed for 3-D surface plots of the biofilms of the other *B. cereus* test isolates.

**Figure 3.**
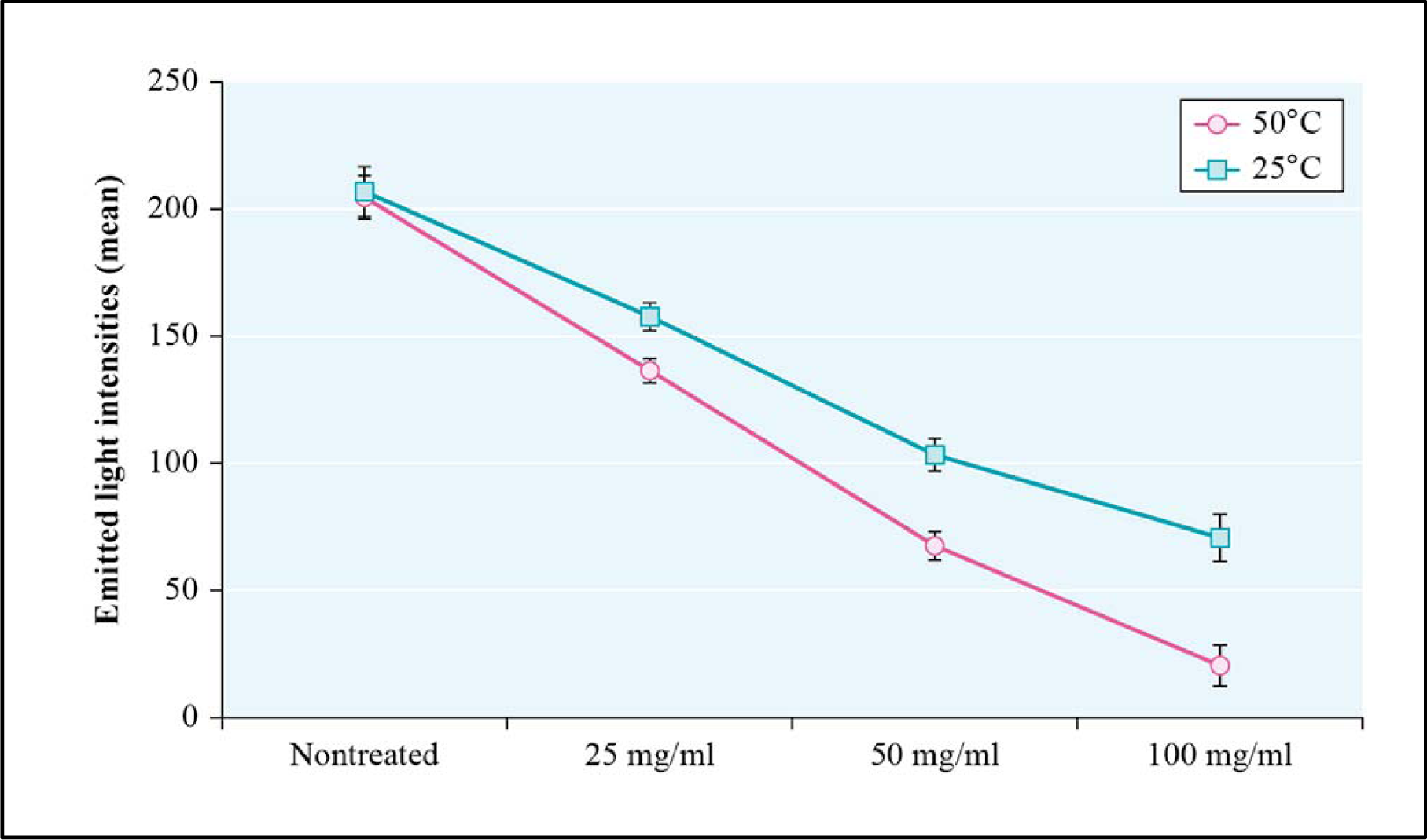
Recorded emitted light intensities in the fluorescence micrographs of BC-MS010 biofilms on SSCs after 15 min immersion in 25, 50, or 100 mg/ml *Pestalotiopsis* sp. culture crude extract at 25°C or 50°C. Error bars indicate the standard deviation from the mean (n=4).

**Figure 4.**
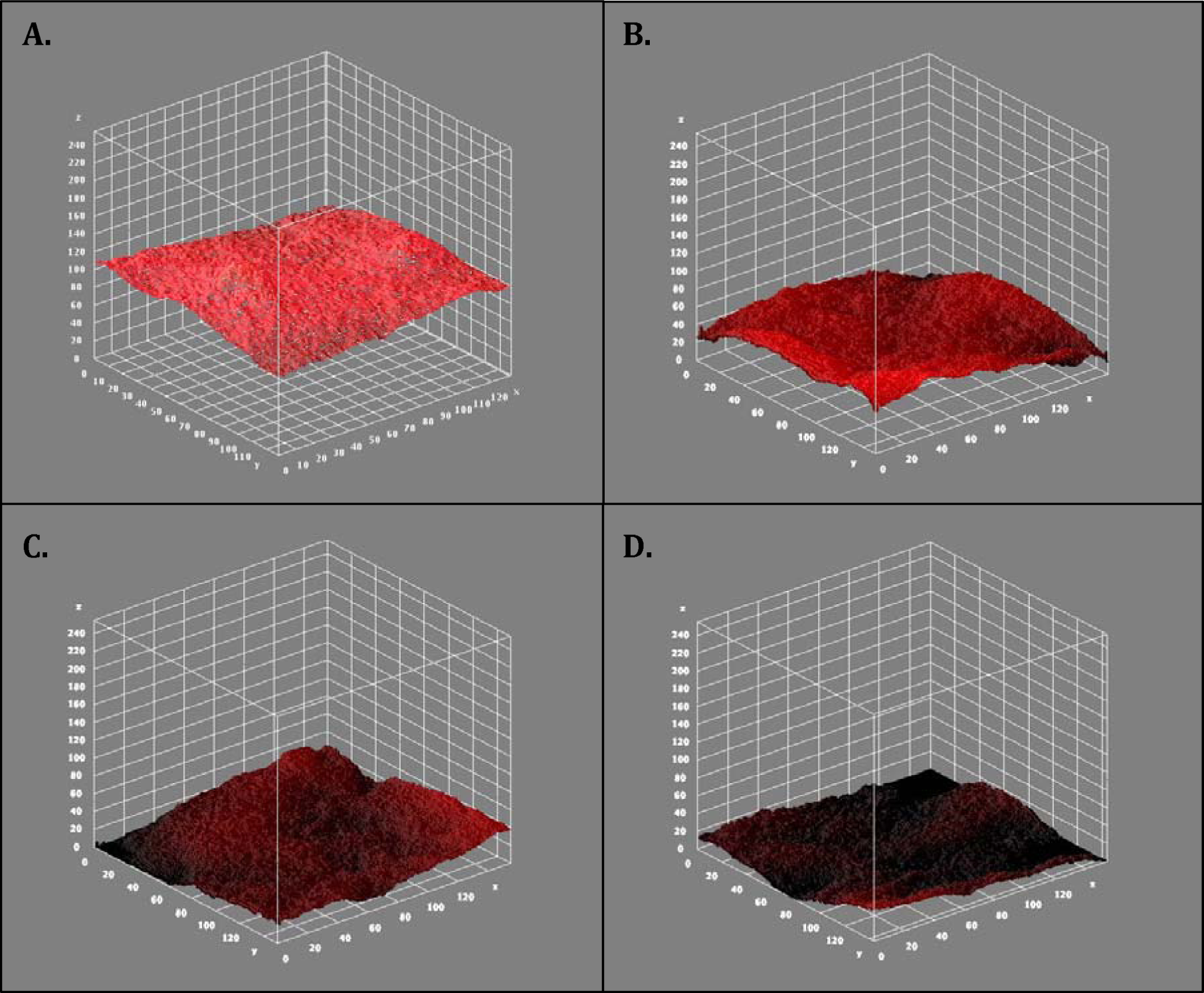
ImageJ-generated interactive 3-D surface plots of the isolate BC-MS010 biofilms on SSCs. Control biofilm (**A**), and biofilms after immersion in (**B**) 25 mg/mL, (**C**) 50 mg/mL, and (**D**) 100 mg/mL *Pestalotiopsis* sp. culture crude extract within 15 min contact time at 50°C.

## 4. Discussion

Microbial biofilms are known to be impervious against most antimicrobial agents, and therefore, biofilms in food production systems resist many kinds of disinfectants. Biofilms can grow on any substrata, such as wood, plastic, and stainless steel surfaces. In this study, the *B. cereus* test isolates were evaluated using the MTP biofilm formation assay, and the isolates that obtained high SBF values were then tested for biofilm formation on SSCs. Also known as microtiter dish assay, MTP biofilm formation assay is a colorimetric assay and is an important tool for studying the early stages of microbial biofilm formation in microtiter plates (O’Toole, 2011). Herein, the attached biofilms on the walls or bottoms of the wells were stained with CV solution. *B. cereus* biofilm production was initially induced within the first 24 hr incubation period at 30°C in diluted TSB in the MTP wells. Previous reports have shown that the switch from the planktonic mode of growth to growth as bacterial biofilms occurs as a response to the availability of nutrients in the environment, with nutrient-depleted medium enhancing biofilm production (Speranza et al., 2011; Thomason et al., 2012; Zhang et al., 2014). The current study supports the findings of Wijman et al. (2007), wherein the authors reported biofilm formation by their *B. cereus* strains within the first 24 hr, followed by subsequent biofilm dispersion over the next 24 h. The authors also concluded that biofilm formation was dependent on strain, incubation time, temperature, and medium. After a total incubation period of 48 hr, we observed that each *B. cereus* isolate formed biofilms at various depths in the MTP wells, and this could be explained by their metabolic preferences and adaptation. For example, the microaerophilic human pathogen *Campylobacter jejuni* develops biofilms rapidly under aerobic condition to survive (Reuter et al., 2010), as this bacterium is known to thrive in anaerobic environments. In general, *B. cereus* is a facultative anaerobe and, therefore, can proliferate with or without oxygen.

The adhesion of *B. cereus* spores to inanimate surfaces is effective due to the hydrophobic and appendage characteristics of the spores (Faille et al., 2010) compared to the vegetative cells. The adhering capacity of spores to stainless steel surfaces is also affected by time, temperature, and pH (Peña et al., 2014). As observed in this study, *B. cereus* spores effectively adhered to milk pre-soiled SSCs, and the biofilms were visible at the ALI after the first 24 hr of incubation under nutrient-depleted and static conditions. By definition, the ALI is the area where microorganisms have the access to both gaseous (i.e., oxygen) and liquid phases, which is advantageous for aerobes and facultative anaerobes (Spiers et al., 2003). However, Wijman et al. (2007) revealed that the pretreatment of SSCs with milk resulted in lower adherence of *B. cereus* spores and subsequently resulted in smaller amounts of biofilm for the majority of the strains tested. In contrast, in this study milk appeared to induce spore attachment to SSCs and eventually lead to biofilm formation by the *B. cereus* test isolates. Such stimulation of spore attachment could result in product contamination if the cleaning and disinfection of stainless steel surfaces fails to remove the milk residues. Additionally, the *B. cereus* biofilms extended below the ALI, where the O_2_ concentration was low, but these biofilms were easily detached by simple rinsing. Therefore, the presence of O_2_ is a vital requirement for biofilm formation by *B. cereus* on any surface and might also contribute to the biofilm thickness, rigidity and, thus, resistance against external factors, such as antibiotics and disinfectants. Additionally, it is clear that *B. cereus* biofilms that form below the ALI can spread freely in food production systems and could increase the magnitude of *B. cereus* contamination. As the biofilms were visibly evident after the first 24 hr incubation period in nutrient-depleted medium, *B. cereus* can form biofilms within a short time period and dwell in less nutritive environments. Furthermore, biofilm formation could occur in food production areas where there are short downtimes between product processing and cleaning, or vice versa.

The crude extract of *Pestalotiopsis* sp. broth culture was subjected to TLC for detection of secondary metabolites before testing against *B. cereus* biofilms on SSCs. As viewed under UV lights, all solvent systems produced spots, suggesting that secondary metabolites were present. However, these compounds remain unknown, and further study is recommended to elucidate their identity. To date, alkaloids, terpenoids, isocoumarin derivatives, coumarins, chromones, quinones, semiquinones, peptides, xanthones, xanthone derivatives, phenols, phenolic acids, and lactones have been reported to be produced by *Pestalotiopsis* spp., making this genus a particularly rich source for bioprospecting compared to other fungal genera (Xu et al. 2010). The crude extract residues have been found to be soluble in methanol and water, but we preferred to use the latter against *B. cereus* biofilms because methanol could dehydrate the biofilms. Based on the results, the *Pestalotiopsis* sp. culture crude extract significantly reduced (p < 0.5) *B. cereus* biofilms on SSCs within 15 min of exposure at 25°C and 50°C. Clear biofilm reduction was observed at 50°C, implying that temperature played an important role in biofilm control. It has been reported that heat shocks between 50°C and 80°C for durations of 1 to 30 min could reduce biofilm CFU concentrations up to 6.6 –fold and are a reasonable approach for biofilm mitigation (O’Toole et al., 2015). Therefore, we conclude that the secondary metabolites extracted from *Pestalotiopsis* sp. have antibiofilm activity against *B. cereus* grown on SSCs. However, it has been noted that 90% of *B. cereus* biofilms are composed of spores (Wijman et al., 2007) and are generally resistant to all kinds of antimicrobial agents and chemicals. Thus, the biofilm cells recovered after immersion in all concentrations of *Pestalotiopsis* sp. culture crude extract probably emerged from spores. We also found that the sterile water washed away the *B. cereus* biofilm cells during the rinsing step after immersion in the fungal crude extract. Additionally, the recorded emitted light from the EtBr attached to the biofilms decreased, based on the RGB values generated by the ImageJ software. Since *Pestalotiopsis* sp. culture crude extract in this study had no antibacterial activity on *B. cereus* vegetative cells, it is inferred that the crude extract only disrupts the biofilms EPS. EPS matrices are primarily composed of proteins (Jiao et al. 2011), DNA (Liu et al., 2011), surfactants (Wilking et al., 2011), glycolipids (Cortes-Sanchez et al., 2012), and ions (Safari et al., 2014; Grumbein et al., 2014). Additionally, the matrix determines the proximate microenvironmental conditions experienced by cells living in the biofilm by affecting porosity, density, water content, charge, sorption properties, hydrophobicity, and mechanical stability (Flemming & Wingender, 2002).

Generally, microbial biofilms have a huge negative impact on food production industries through product contamination and the spread of foodborne illness. In this study, the crude extract from *Pestalotiopsis* sp. broth culture demonstrated antibiofilm activity against *B. cereus* grown on stainless steel surfaces. Stainless steel is the most common type of steel alloy used in most parts of food production systems because of its corrosion resistance and low maintenance, but it is a good attachment site for most microbial food pathogens, including *B. cereus*. To the best of our knowledge, this is the first report of secondary metabolites from the crude extract of *Pestalotiopsis* sp. being tested against bacterial biofilms. Additionally, this study offers evidence that fungal plant endophytes, such as the *Pestalotiopsis* spp. and related genera, could be a source of natural products that could function as alternatives to existing antimicrobial agents that are no longer effective due to microbial resistance. Hence, the current study provides additional and promising results that could give further perspective to biofilm control and eradication in food manufacturing systems.

## Declarations

### Author contribution statement

Rener De Jesus and Gina Dedeles performed the experiments, analyzed and interpreted the data, and wrote the paper.

### Additional information

No additional information is available.

## Acknowledgment

We thank Dr. Irineo J. Dogma, Jr. for confirming the conidial morphology and characteristics of our *Pestalotiopsis* sp. isolate.

## References

Ash, C., Farrow, J. A., Dorsch, M., Stackenbrandt, E., Collins, M. D., 1991. Comparative analysis of *Bacillus anthracis, Bacillus cereus*, and related species on the basis of reverse transcriptase of 16S rRNA. International Journal of Systematic Bacteriology, 41, 343–346. doi: 10.1099/00207713-41-3-343

Bai, Z., Lin, X., Wang, Y., Wang, J., Zhou, X., Yang, B., Liu, J., Yang, X., Wang, Yi., Liu, W., 2014. New phenyl derivatives from endophytic fungus *Aspergillus flavipes* AIL8 derived of mangrove plant *Acanthus ilicifolius*. Fitoterapia, 95, 194–202. doi: 10.1016/j.fitote.2014.03.021

Bayoumi, M., Kal., R., Abd El Aal, S., Awad, E., 2012. Assessment of a regulatory sanitization process in Egyptian dairy plants in regard to the adherence of some food-borne pathogens and their biofilms. International Journal of Food Microbiology, 158, pp.225–231. doi: 10.1016/j.ijfoodmicro.2012.07.021

Beecher, D. J., MacMillan, J. D., 1991. Characterization of the components of hemolysin BL from *B. cereus*. Infection and Immunity, p. 1778–1784, Vol. 59, No. 5.

Brézillon, C., Haustant, M., Dupke, S., Corre, J-P., Lander, A., Franz, T., Monot, M., Couture-Tosi, E., Jouvion, G., Leendertz, F., Grunow, R., Mock, M., Klee, S., Goossens, P., 2015. Capsules, toxins and ATxa as virulence factors of emerging *B. cereus* biovar *anthracis*. PLOS Neglected Tropical Diseases 9(4): e0003746. doi: 10.1371/journal.pntd.0003455

Chang, H., Lee, J., Han, B., Kwak, T., Kum, J., 2011. Prevalence of the levels of *B. cereus* in fried rice dishes and its exposure assessment from Chinese-style restaurants. Food Science and Biotechnology, 20:1351. doi: 10.1007/s10068-011-0186-3

Chen, S., Qian, L., Shi, B., 2007. Purification and properties of enantioselective lipase from a newly isolated *Bacillus cereus* C71. Process Biochemistry, 42, 988–994. doi: 10.1016/j.procbio.2007.03.010

Cortes-Sanchez, A., Hernandez-Sanchez, H., Jaramillo-Flores, M. E., 2012. Biological activity of glycolipids produced by microorganisms: new trends and possible therapeutic alternatives. Microbiological Research, 168, 22–32. doi: 10.1016/j.micres.2012.07.002

Donlan, R., Costerton, J., 2002. Biofilms: survival mechanisms of clinically relevant microorganisms. Clinical Microbiology Reviews. Vo. 15, No. 2, pp. 167–193. doi: 10.1128/CMR.15.2.167-193.2002

Elhariry, H. M., 2011. Attachment strength and biofilm forming ability of *B. cereus* on green-leafy vegetables: cabbage and lettuce. Food Microbiology, 28, 1266–1274. doi: 10.1016/j.fm.2011.05.004

Eneroth, A., Svensson, B., Molin, G., Christiansson, A., 2001. Contamination of pasteurized milk by *B. cereus* in filling machines. Journal of Dairy Research. Vol. 76, Issue 2, pp. 189–196. doi: 10.1017/S002202990100485X

Faille, C., Fontaines, F., Bénézech, T., 2001. Potential occurrence of adhering living Bacillus spores in milk product processing lines. Journal of Applied Microbiology, 90, 892–900. doi: 10.1046/j.1365-2672.2001.01321.x

Faille, C., Lequette, Y., Ronse, A., Slomianny, C., Garénaux, Guerardel, Y., 2010. Morphology and physic-chemical properties of Bacillus spores surrounded or not with an exosporium consequences on their ability to adhere to stainless steel. International Journal of Food Microbiology, 143, 125–135. doi: 10.1016/j.ijfoodmicro.2010.07.038

Flemming, H. C., and Wingender, J. in Encyclopedia of Environmental Microbiology (ed. Bitton, G.) 1223–1231 (Wiley, New York, 2002).

Grumbein, S., Opitz, M., Lieleg, O., 2014. Selected metal ions protect *Bacillus subtilis* biofilms from erosion. Metallomics, 6, 1441. doi: 10.1039/c4mt00049h

Hwang, I. H., Swenson, D., Gloer, J., Wicklow, D., 2015. Pestaloporonins: caryophylle-derived sesquiterpenoids from a fungicolous isolate of *Pestalotiopsis* sp. Org. Lett., 17. 4284–4287. doi: 10.1021/acs.orglett.5b02080

Jiao, Y., D’haesseleer, P., Dill, B. D., Shah, M., VerBerkamoes, N. C., Hettich, R. L., Banfield, J., Thelen, M., 2011. Identification of biofilm matrix associated proteins from an acid mine drainage microbial community. Applied Environmental Microbiology. doi: 10.1128/AEM.03005-10

Kim, S., Young, K., Rhee, I-K., 2001. Purification and characterization of a novel extracellular protease from *B. cereus* KCTC 3674. Archives of Microbiology, Vol. 175, Issue 6, pp. 458–461. doi: 10.1007/s002030100282

Li, X., Guo, Z., Deng, Z., Yang, J., Zou, K., 2015. A new α-pyrone derivative from endophytic fungus *Pestalotiopsis microspora*. Records of Natural of Products, 9:4, 503–508.

Liu, H-H., Yang, Y-R., Li, Y., Xie, Z-X., Shen, P., 2011. Involvement of DNA in biofilm formation: features and mechanism of DNA adsorption to bacterial surfaces. The Journal of Bioscience and Medicine, 6. doi: 10.5780/jbm2011.6

Lund, T., and Granum, P. E., 1996. Characterization of a non-haemolytic enterotoxin complex from *B. cereus* isolated after a foodborne outbreak. FEMS Microbiology Letters, 141, 151–156. doi: 10.1111/j.1574-6968.1996.tb08377.x

Martinez-Medina, M., Naves, P., Blanco, J., Aldeguer, X., Blanco, J., Blanco M., Ponte, C., Soriano, F., Darfueille-Michaud, A., Garcia-Gil, L.J., 2009. Biofilm formation as a novel phenotypic feature of adherent-invasive *Escherichia coli* (AIEC). BMC Microbiology, 9:202. doi: 10.1186/1471-2180-9-202

Møretrø, T., Schirmer, B., Heir, E., Fagerlund, A., Hjemli, P., Langsrud, S., 2017. Tolerance to quaternary ammonium compounds disinfectants may enhance growth of *L. monocytogenes* in the food industry. International Journal of Food Microbiology, 241, 215–224. doi: 10.1016/j.ijfoodmicro.2016.10.025

Niu, C., and Gilbert, E. S., 2004. Colorimetric method for identifying plant essential oil components that affect biofilm formation and structure. Applied and Environmental Microbiology, 70, 6951–6956. doi: 10.1128/AEM.70.12.6951-6956.2004

O’Toole, A., Ricker, E., and Nuxoll, E., 2015. Thermal mitigation of *Pseudomonas aeruginosa* biofilms, Biofouling, 31 (8): 665–675. doi: 10.1080/08927014.2015.1083985

O’Toole, G., 2011. Microtiter Dish Biofilm Formation Assay. J. Vis Exp.; (47):2437. doi: 10.3791/2437

Peña, W., de Andrade, N., Soares, N., Alvarenga, V., Junior, S., Granato, D., Zuniga, A., Sant’Ana, A., 2014. Modelling *B. cereus* adhesion on stainless steel surface as affected by temperature, pH, and time. International Dairy Journal, 34, 153–158. doi: 10.1016/j.idairyj.2013.08.006

Rajkovic, A., Uyttendaele, M., Dierick, K., Samapundo, S., Botteldoorn, N., Mahillon, J., Heyndrickx, M., 2008. Risk profile of the *B. cereus* group implicated in food poisoning. Report for the Superior Health Council Belgium.

Reuter, M., Mallett, A., Pearson, B., van Vliet, A., 2010. Biofilm formation by *Campylobacter jejuni* is increased under aerobic conditions. Applied and Environmental Microbiology, Vol. 76, No. 7, p. 2122–2128. doi: 10.1128/AEM.01878-09

Safari, A., Habimana, O., Allen, A., Casey, E., 2014. The significance of calcium ions on *Pseudomonas fluorescens* biofilms-a structural and mechanical study. Biofouling, 30:7, 859–869. doi: 10.1080/08927014.2014.938648

Schulz, B., Wanke, U., Draeger, S. and Aust, H. J., 1993. Endophytes from herbaceous plants and shrubs, effectiveness of surface sterilization methods. Mycological Research, 97, 1447–1450. doi: 10.1016/S0953-7562(09)80215-3

Sharma, D., Pramanik, A., Agrawal, P., 2016. Evaluation of bioactive secondary metabolites from endophytic fungus *Pestalotiopsis* BAB-5510 isolated from leaves of *Cupressus torulosa* D. Don. Biotech, 6:210. doi: 10.1007/s13205-016-0518-3

Speranza, B., Corbo, M. R., Sinigaglia, M., 2011. Effects of nutritional and environmental conditions on *Salmonella* sp. biofilm formation. Journal of Food Science, Volume 76, Issue 1. doi: 10.1111/j.1750-3841.2010.01936.x

Spiers, A. J., Bohannon, J., Gehrig, S. M., Rainey, P. B., 2003. Biofilm formation at the air-liquid interfaces by the *P. fluorescens* SBW25 wrinkly spreader requires an acetylated form of cellulose. Molecular Microbiology, 50, 15–27. doi: 10.1046/j.1365-2958.2003.03670.x

Stenfors Arnesen, L. P., Fagerlund, A., Granum, P. E., 2008. From soil to gut: *B. cereus* and its food poisoning toxins. FEMS Microbiological Reviews 32:579–606. doi: 10.1111/j.1574-6976.2008.00112.x

Svensson, B., Ekelund, K., Ogura, H., Christiansson, A., 2004. Characterisation of *B. cereus* isolated from milk silo tanks at eight different dairy plants. International Dairy Journal, 14, 17–27. doi: 10.1016/S0958-6946(03)00152-3

Tewari, A., Singh, S. P., Singh, R., 2015. Incidence and enterotoxigenic profile of *B. cereus* in meat and meat products of Uttarakhand, India. Journal of Food Science and Technology, Vol. 52, Issue 3, pp 1796–1801. doi: 10.1007/s13197-013-1162-0

Thomason, M. K., Fontaine, F., De Lay, N., Storz, G., 2012. A small RNA that regulates motility and biofilm formation in response to changes in nutrient availability in *E. coli*. Molecular Microbiology, 84 (1), 17–25. doi: 10.1111/j.1365-2958.2012.07965.x

Wijman, J., de Leeuw, P., Moezelaar, R., Zwietering, M., Abee, T., 2007. Air-liquid interface biofilms of *Bacillus cereus* L formation, sporulation and dispersion. Applied and Environmental Microbiology, Vol. 73, No. 5, p 1481–1488. doi: 10.1128/AEM.01781-06

Wilking, J. N., Angelini, T. E., Seminara, A., Brenner, M. P., Weitz, D. A., 2011. Biofilms as complex fluids. MRS Bulletin, 36, 385–391. doi: 10.1557/mrs.2011.71

Xu, J., Ebada, S., Proksch, P., 2010. *Pestalotiopsis* as highly creative genus: chemistry and bioactivity of secondary metabolites. Fungal Diversity, Vol. 44, Issue I., pp. 15–31. doi: 10.1007/s13225-010-0055-z

Yang, Y. Mikš-Krajnik, M., Zheng, Q., Lee, S., Lee, S., Yuk, H., 2016. Biofilm formation of *Salmonella enteritidis* under food-related environmental stress conditions and its subsequent resistance to chlorine treatment. Food Microbiology, 34, 98–105. doi: 10.1016/j.fm.2015.10.010

Zhang, W., Seminara, A., Suaris, M., Brenner, M., Weitz, D., Angelini, T., 2014. Nutrient depletion in *Bacillus subtilis* biofilms triggers matrix production. New Journal of Physics 16, 015028. doi: 10.1088/1367-2630/16/1/015028

Zhang, Z., Feng, L., Xu, H., Liu, C., Shah, N., Wei, H., 2016. Detection of viable enterotoxin-producing *B. cereus* and analysis of toxigenicity from ready-to-eat foods and infant formula milk powder by multiplex PCR. Journal of Dairy Science, Vol.99, Issue 2, pp. 1047–1055. doi: 10.3168/jds.2015-10147

Zhao, J., Mou, Y., Shan, T., Li, Y., Zhou, L., Wang, M., Wang, J., 2010. Antimicrobial metabolites from the endophytic fungus *Pichia guilliermondii* isolated from *Paris polyphylla* var. *yunnanensis*. Molecules 2010, 15, 7961–7970; doi: 10.3390/molecules15117961

Zin, W., Buttachon, S., Dethoup, T., Pereira, J., Gales, L., Inácio, A., Costa, P., Lee, M., Sekeroglu, N., Silva, A., Pinto, M., Kijjoa, A., 2017. Antibacterial and antibiofilm activities of the metabolites isolated from the culture of the mangrove-derived endophytic fungus *Eurotium chevalieri* KUFA 0006. Phytochemistry, 141, 86–97. doi: 10.1016/j.phytochem.2017.05.015

